# Courtship vocalizations in male ducks: spectral composition and resonance of the syringeal bulla

**DOI:** 10.1101/2025.01.13.632779

**Authors:** Darcy Mishkind, Michael Lester, Clifford J. Tabin, Franz Goller

## Abstract

Ducks display a unique and dramatic sexual dimorphism in their vocal organ, the syrinx. Males have a left-sided bulla that is not present in females and that has been long hypothesized to play a role in courtship vocalizations, though this connection has never been tested. The large, hollow morphology of the bulla and its proximity to the sound-producing vocal folds introduce the possibility that it may work as a Helmholtz resonator, which makes it possible to predict the resonance frequencies enhanced by this structure. We find that during early ontogeny, the distribution of energy across the harmonic spectrum of contact calls is not different between males and females. We then used micro-CT scans of duck syrinxes to estimate resonance frequencies of the bullae and compared these to spectral features of their vocalizations. This comparison overall supports the idea that the bulla resonance may specifically enhance aspects of courtship vocalizations, especially in species that have a tonal courtship whistle. We also see potential influence in non-courtship vocalizations, which could be explored further with a greater understanding of the input of other vocal tract features that influence vocalization. We observed that bulla size is positively correlated with bird body mass with the largest exception in the Common Eider, which had a small bulla for its body mass and for which we saw no evidence of bulla-input to its vocalizations. This study found support for the long-held hypothesis that the adult male duck bulla influences resonance frequencies, in particular in courtship vocalizations.

## Introduction

Birds generate sound with a unique, avian specific vocal organ located at the tracheobronchial junction, called the syrinx. The syrinx consists of modified cartilage elements of the trachea and bronchi, which house the sound generating vibratory tissue (King, 1989; Goller and Larsen, 1997; Elemans et al., 2015). Depending on the specific location of these vocal folds (often also called membranes or labia) in the tracheal or bronchial parts of the syrinx, birds can have one, two or three sound sources (King, 1989; Goller, 2016; Garcia et al., 2017). It is likely that a syrinx with two bronchial sound generators represents the original syrinx morphology (Clarke et al., 2016; Longtine et al., 2024), and this vocal structure is found in many avian orders, including the basal galloanserines. The vocal folds are suspended in a framework of ossified cartilages, which are often modified from typical bronchial or tracheal elements. High levels of variation in these structures have inspired scientists early on to use the syrinx as a taxonomic characteristic for classifying birds (Fürbringer and Fürbringer 1888; Gadow and Selenka 1891; Wunderlich 1884). While modern taxonomic methods have largely replaced these early classification systems, morphological characteristics of the syringeal skeletal elements still inspire research on the evolution of this unique vocal organ.

The morphological variety of the syrinx and vibratory tissues contributes to the large acoustic range of avian vocalizations (Gaunt, 1983; Riede and Goller, 2014; Goller, 2022). In many avian taxa, vocal behavior also differs between males and females. However, we know relatively little to what degree these differences are rooted in sexual dimorphism of the vocal apparatus. Although the syringeal structures of male zebra finches are larger than those of females (Düring et al., 2013), analysis of differences in volume, fiber type composition and contraction dynamics of syringeal musculature indicate that neuromuscular control predominantly accounts for different acoustic output (Wade and Buhlman, 2000; Elemans et al., 2008; Christensen et al., 2017).

The most striking sexual dimorphism in syringeal structure can be found in ducks. The syrinx of male ducks exhibits a bulb-like, air-filled outgrowth of the left syringeal skeleton, referred to as a bulla (Rüppell, 1933; Johnsgard, 1961; King, 1989; Frank et al., 2007). In Mallards (*Anas platyrhynchos*), the bulla is present at hatching, but sexual dimorphism of the syringeal labia and membranes is only expressed later (>45 days post hatching) (Gottlieb and Vandenbergh, 1968; Warner, 1971; Lockner and Youngren, 1976). While duckling calls do not appear sexually dimorphic, adult male and female ducks of many species have distinct vocal repertoires (Abraham, 1974; Lockner and Murrish, 1975). In particular, in many species males produce distinct courtship vocalizations (Johnsgard, 1971). It is tempting therefore to associate the presence of a bulla with the generation of courtship vocalizations (Johnsgard, 1961, 1971). While several possible roles of the bulla in the production or modification of vocalizations in the respective vocal repertoires have been proposed (Johnsgard, 1971; Lockner and Youngren, 1976), they have, to our knowledge, not been thoroughly tested.

How could the presence of a bulla influence vocal production? Ducks possess two sound generators (membranes or labia), and the bulla opening is situated close to the left labial pair. In Mallards the labia and attached medial tympaniform membranes are also sexually dimorphic, being small in females and thick in males (Warner, 1971). It is likely that oscillations of the labia are the primary sound source, although there is some disagreement in the literature (discussed in King 1989). Sound production in male Mallards appears to be linked to the presence of the bulla, such that the right pair of labia is not engaged (Lockner and Youngren, 1976).

Irrespective of the precise oscillatory mechanism, the generated sound is modified by the resonance properties of the upper vocal tract structures. The tracheal tube, oropharyngeal-esophageal cavity (OEC), glottal opening, mouth, tongue and bill provide static and dynamically adjustable mechanisms for upper vocal tract filtering and thus contribute to the spectral properties of emitted sound (Riede et al., 2006; Ohms et al., 2010, 2012; Beckers, 2013; Kazemi et al., 2023). In male ducks, the bulla is also predicted to contribute specific resonance properties (Abs, 1969, 1970a, 1970b; Warner, 1971; Abraham, 1974). Its morphology and proximity to the sound source suggest that it may act as a Helmholtz resonator. Its geometry should therefore allow predictions about its resonance properties and, thus, which spectral features of vocalizations are enhanced. The presence of a bulla only in males opens an intriguing opportunity to ask how this specific feature contributes to vocalization.

While its function is untested, an impressive diversity of bulla morphology has been described (Johnsgard, 1961). The wide array of sizes has produced a ‘natural experiment’: As the size varies, so too does the frequency a given bulla is predicted to amplify. Taking advantage of this, we compared the vocalizations of different species to the frequencies we would predict based on their bullae morphology, to test whether there is evidence that the bulla is used for male-specific sound production. Here, we have made used of a wide collection of duck species found in a museum collection to produce µCT scans of duck syrinxes. We hypothesized that the bulla acts as a resonance chamber. On this basis we used the scans to model predicted bulla-produced frequencies and asked how these compare to the recorded vocal repertoires of a given species. Due to the strong sexual dimorphism displayed in this trait, we expected it to play a role in the spectral composition of duckling calls and male-specific adult vocalizations.

## Results

We began by exploring the developmental progression of duckling contact calls in order to evaluate if the presence of a bulla in males affects the spectral energy distribution. The frequency contour of contact calls is hat-like with rising and declining frequency at the onset and offset (Fig. 1A). The fundamental frequencies have the highest relative amplitude (dB) and harmonics are typically not emphasized (Figs 1,2). Energy distributions across the measured time scale were very similar for males and females (Fig. 2A). The only exception is around age 32 days post hatching, where the mean F2 amplitude in males is distinctly higher (approx. 20 dB) than that at all other time points and notably higher than that of females (Fig. 2A). At this time point the FF of the four male ducklings ranged from 2.2–2.7 kHz (Fig. 2B), thus providing a range of the F2 from 4.4–5.4 kHz.

**Figure 1.**
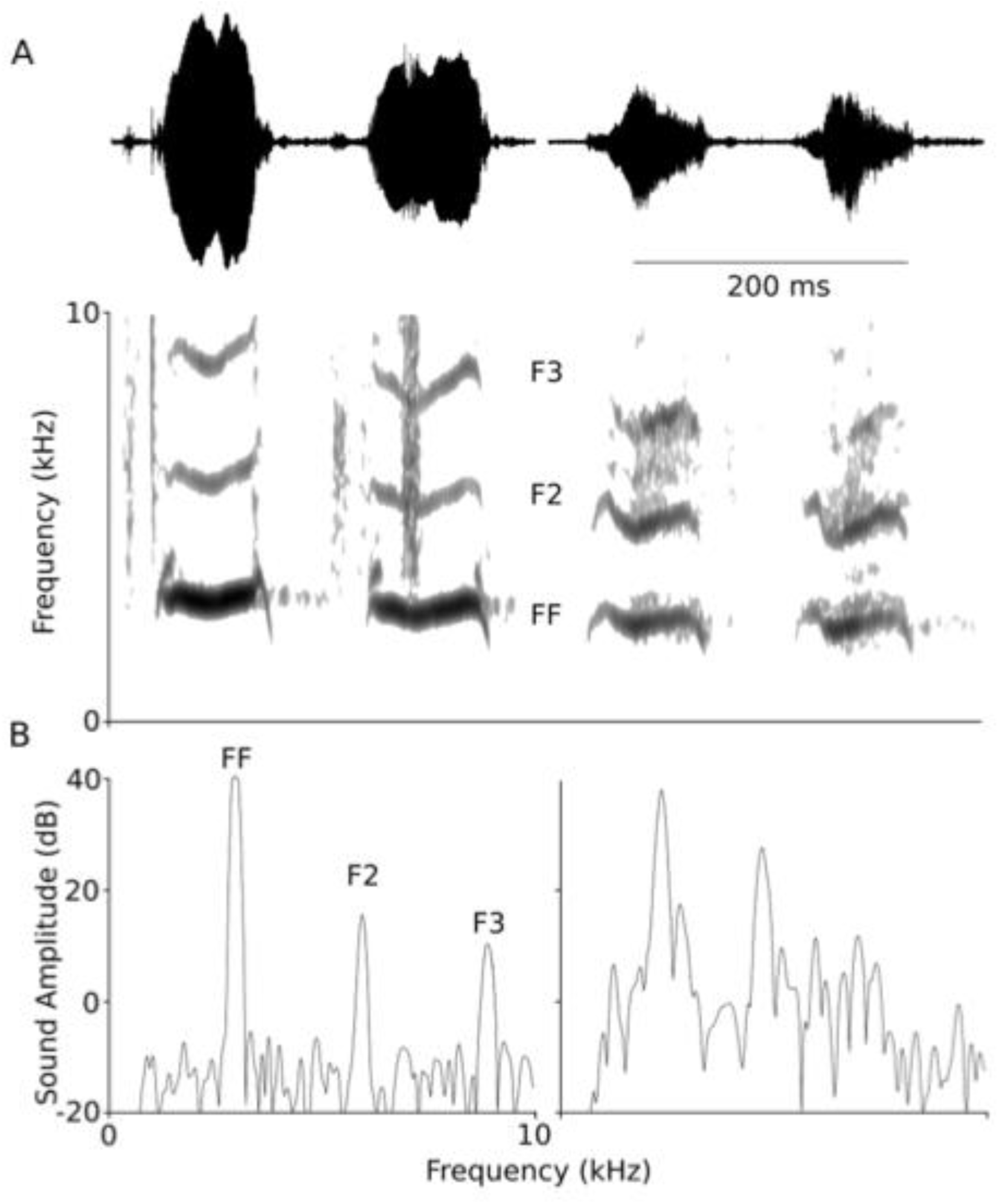
Male duckling contact calls. (A) Contact call examples (shown as oscillogram and spectrogram) from a male duckling at 11 and 32 days of age illustrate the decrease in fundamental frequency (FF) and the harmonic content (F2, second harmonic, F3, third harmonic). (B) Power spectra of the center region of the first calls at each age show the increased emphasis of the second harmonic at the older age (right).

**Figure 2.**
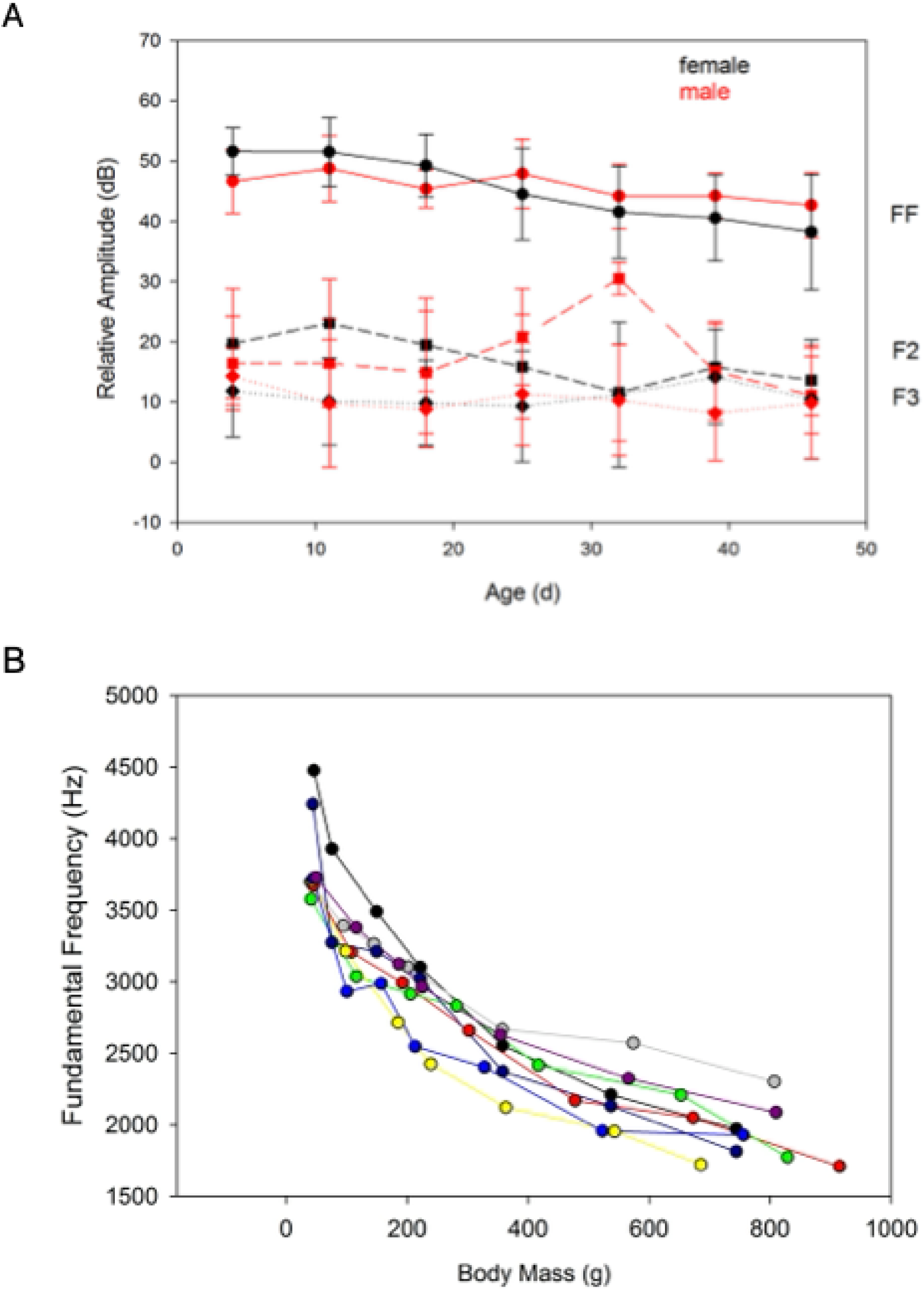
Male and female duck contact calls show similar trends in amplitude and fundamental frequency over developmental time. (A) Male and female contact calls show similar relative amplitudes (means ±1 s.d.) of fundamental frequency (FF, circles and solid lines), second (F2, squares and stippled lines) and third (F3, diamonds and dotted lines) over the age range. The exception is a more pronounced second harmonic at 32 days in male calls. Call amplitude declined slightly as ducklings tended to give softer contact calls with increasing age. (B) The fundamental frequency (means) of contact calls declines as ducklings grow. Different colors are used for the data of different individuals.

We then turned to the potential function of the duck bulla. We took advantage of the Harvard Museum of Comparative Zoology ornithology collections to study 17 species of duck with intact syrinxes. We used µCT scans to establish the dimensions of the bullae (Fig. 3; Table 1). Three of the species had a smaller right bulla in addition to the left bulla: the Common Goldeneye, Red-Breasted Merganser, and Common Shelduck (Table 1). These measurements were then used to model each bulla as a Helmholtz resonator and predict resonance frequencies (Table 1).

**Figure 3.**
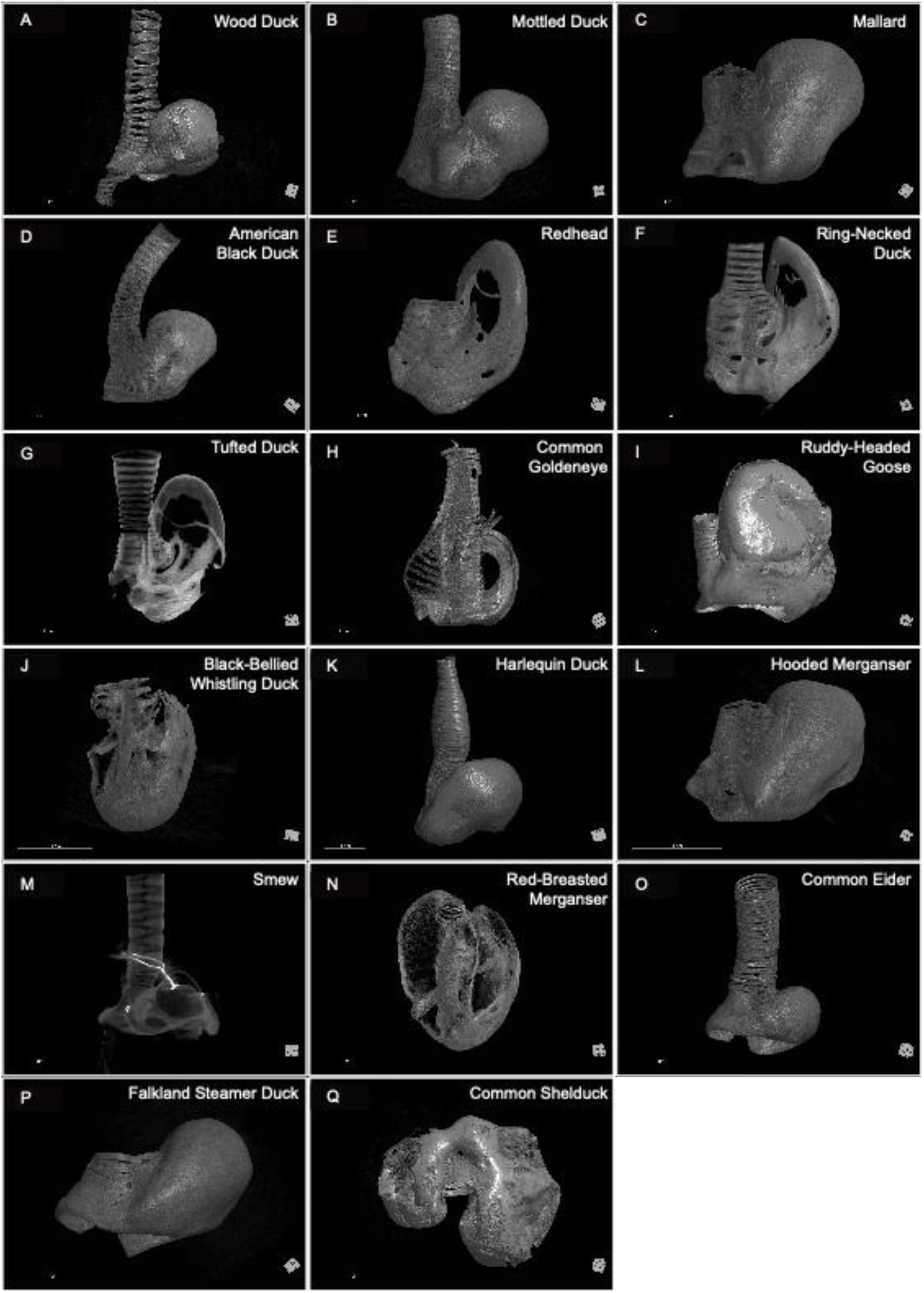
Two-dimensional renderings of µCT-scanned syrinxes. Ventral view. (A–Q), Species and sample indices as follows: Wood Duck (*Aix sponsa*), 341876; Mottled Duck (*Anas fulvigula*), 342070; Mallard (*Anas platyrhynchos*), 347156; American Black Duck (*Anas rubripes*), 335525; Redhead (*Aythya americana*), 346918; Ring-Necked Duck (*Aythya collaris*), 341918; Tufted Duck (*Aythya fuligula*), 340385; Common Goldeneye (*Bucephala clangula*), 347155; Ruddy-Headed Goose (*Chloephaga rubidiceps*), 343209; Black-Bellied Whistling Duck (*Dendrocygna autumnalis*), 340273; Harlequin Duck (*Histrionicus histrionicus*), 336960; Hooded Merganser (*Lophodytes cucullatus*), 342699; Smew (*Mergus albellus*), 340383; Red-Breasted Merganser (*Mergus serrator*), 341905; Common Eider (*Somateria mollissima*), 337334; Falkland Steamer Duck (*Tachyeres brachypterus*), 342206; and Common Shelduck (*Tadorna tadorna*), 347540. Mallard is reproduced for comparison (Mishkind et al., 2024). Scale bar is 1cm.

**Table 1.** µCT-scanned duck species, sample measurements, and calculated resonance frequencies.

| Scientific Name | Common Name | Catalog Number | Side | Bulla Volume (mm<sup>3</sup>) | Opening Diameter (mm) | Neck Length (mm) | Resonance Frequency (Hz) |
| --- | --- | --- | --- | --- | --- | --- | --- |
| *Aix sponsa* | Wood Duck | MCZ:Orn:341876 | Left | 1240.86 | 5.75 | 0.86 | 3405.92 |
| *Aix sponsa* | Wood Duck | MCZ:Orn:347504 | Left | 1266.41 | 4.96 | 0.8 | 3111.78 |
| *Anas fulvigula* | Mottled Duck | MCZ:Orn:342070 | Left | 3197.14 | 6.17 | 1.14 | 2157.74 |
| *Anas platyrhynchos* | Mallard | MCZ:Orn:347156 | Left | 3549.71 | 9.7 | 0.54 | 2757.61 |
| *Anas rubripes* | American Black Duck | MCZ:Orn:335525 | Left | 3508.36 | 8.01 | 1.54 | 2337.92 |
| *Anas rubripes* | American Black Duck | MCZ:Orn:346547 | Left | 3678.73 | 9.44 | 1.96 | 2459.33 |
| *Aythya americana* | Redhead | MCZ:Orn:346918 | Left | 2095.85 | 6.03 | 0.87 | 2691.36 |
| *Aythya collaris* | Ring-Necked Duck | MCZ:Orn:341918 | Left | 2015.66 | 8.27 | 1.88 | 3079.34 |
| *Aythya fuligula* | Tufted Duck | MCZ:Orn:340385 | Left | 1997.09 | 8.22 | 0.86 | 3290.00 |
| *Bucephala clangula* | Common Goldeneye | MCZ:Orn:347155 | Left | 5233.93 | 4.08 | 3.8 | 1030.91 |
| *Bucephala clangula* | Common Goldeneye | MCZ:Orn:347155 | Right | 1308.1 | 6.28 | 2.05 | 3179.06 |
| *Chloephaga rubidiceps* | Ruddy-Headed Goose | MCZ:Orn:343209 | Left | 6744.18 | 6.46 | 1.54 | 1479.81 |
| *Dendrocygna autumnalis* | Black-Bellied Whistling Duck | MCZ:Orn:340273 | Left | 1,055.14 | 2.96 | 1.07 | 2392.71 |
| *Histrionicus histrionicus* | Harlequin Duck | MCZ:Orn:336269 | Left | 3798.3 | 4.24 | 0.75 | 1647.77 |
| *Histrionicus histrionicus* | Harlequin Duck | MCZ:Orn:336960 | Left | 4368.48 | 4.56 | 1.33 | 1506.11 |
| *Lophodytes cucullatus* | Hooded Merganser | MCZ:Orn:342699 | Left | 1242.79 | 5.24 | 0.84 | 3230.34 |
| *Mergus albellus* | Smew | MCZ:Orn:340383 | Left | 2214.95 | 5.39 | 1.87 | 2242.77 |
| *Mergus serrator* | Red-Breasted Merganser | MCZ:Orn:341891 | Left | 6969.61 | 6.98 | 1.17 | 1568.31 |
| *Mergus serrator* | Red-Breasted Merganser | MCZ:Orn:341891 | Right | 4734.47 | 5.93 | 1.18 | 1725.71 |
| *Mergus serrator* | Red-Breasted Merganser | MCZ:Orn:341905 | Left | 6640.54 | 7.52 | 1.36 | 1656.23 |
| *Mergus serrator* | Red-Breasted Merganser | MCZ:Orn:341905 | Right | 3956.18 | 6.31 | 1.26 | 1946.70 |
| *Mergus serrator* | Red-Breasted Merganser | MCZ:Orn:342835 | Left | 8431.59 | 8.9 | 1.37 | 1621.77 |
| *Mergus serrator* | Red-Breasted Merganser | MCZ:Orn:342835 | Right | 5280.2 | 8.9 | 1.84 | 1994.08 |
| *Somateria mollissima* | Common Eider | MCZ:Orn:336858 | Left | 1361.12 | 6.07 | 1.43 | 3197.39 |
| *Somateria mollissima* | Common Eider | MCZ:Orn:337334 | Left | 1439.66 | 7.14 | 1.49 | 3417.18 |
| *Somateria mollissima* | Common Eider | MCZ:Orn:342589 | Left | 1377.45 | 6.51 | 1.2 | 3377.43 |
| *Somateria mollissima* | Common Eider | MCZ:Orn:342590 | Left | 1409.08 | 7.62 | 1.15 | 3676.69 |
| *Somateria mollissima* | Common Eider | MCZ:Orn:342591 | Left | 1362.43 | 8.27 | 2.11 | 3695.10 |
| *Tachyeres brachypterus* | Falkland Steamer Duck | MCZ:Orn:342206 | Left | 3296.15 | 8.22 | 1.37 | 2476.04 |
| *Tadorna tadorna* | Common Shelduck | MCZ:Orn:347540 | Left | 3694.86 | 8.34 | 1.36 | 2360.11 |
| *Tadorna tadorna* | Common Shelduck | MCZ:Orn:347540 | Right | 1772.15 | 8.32 | 1.22 | 3433.64 |

The bulla frequency predictions are based on three key measurements: the bulla volume, the opening diameter, and the neck length of the opening. It is difficult to get precise estimates for the latter two variables, and the opening diameter may be subject to dynamic adjustment. For these reasons, we established how much our estimates for resonance frequency are altered by a 1 mm increase and decrease in these parameters. We used two example species. The change in opening diameter resulted in a maximum average change of 13% for Red-Breasted Merganser and 14% for Common Eider. The change in neck length resulted in a maximum average change in length of 77% for Red-Breasted Merganser and 71% for Common Eider. From this we established that we could expect at maximum a roughly 500–1000 Hz range in resonance frequencies from a combined change in opening diameter and neck length. Opening diameter and neck length each contributed approximately equally to the change in resonance frequency (Fig. 4).

**Figure 4.**
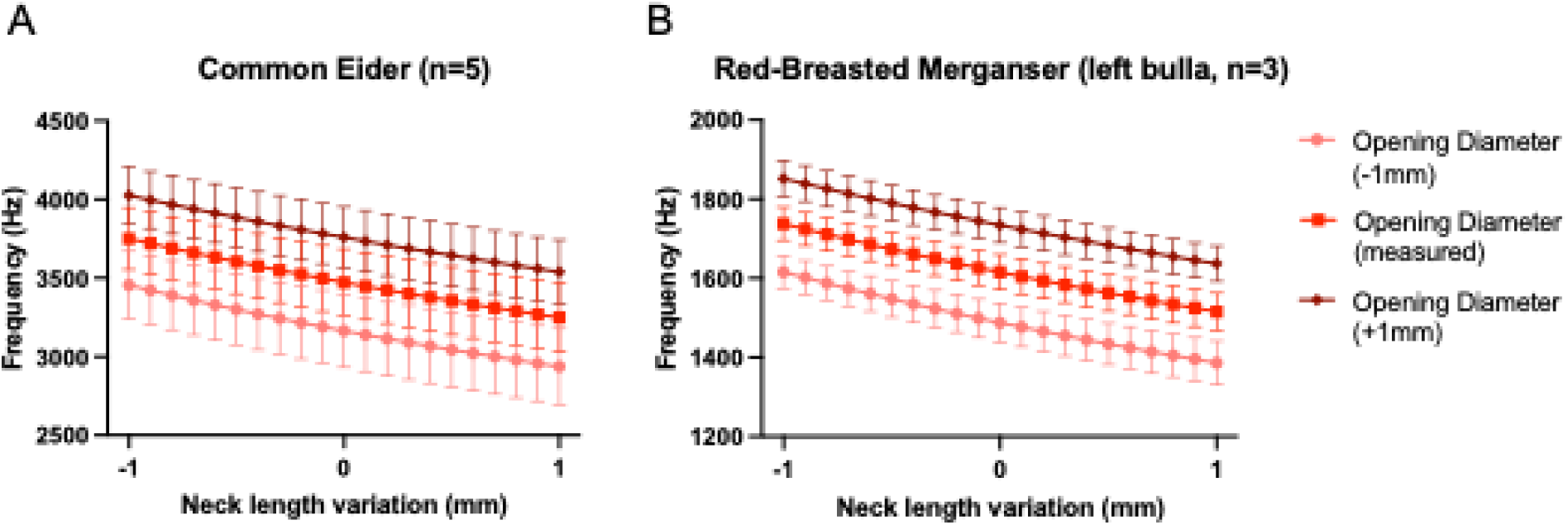
Model of influence of opening diameter and neck length measurements on predicted resonance frequencies. Opening diameter was calculated for 1 mm smaller (pink circles) than measured (red squares) to 1 mm larger than measured (dark red diamonds). Neck length varied from 1 mm smaller than measured to 1 mm larger than measured. (A) Common Eider. (B) Red-Breasted Merganser. Error bars show ± 1 s.d.

We then compared the spectral energy distributions of different call types and compared them to the predicted resonance frequencies. Fig. 5 shows examples for call types from 3 species and the corresponding power spectra. The spectral energy distribution was then plotted and compared to the predicted bulla frequencies (Fig. 6, Supplemental Figs 1, 2). For 10 out of 16 species, the predicted resonance frequency of the bulla is within 500 Hz of a peak in the power spectrum, and 13 out of 16 species were within 1000 Hz (Fig. 6). The courtship calls of the Mallard and closely related species are tonal, whistle-like vocalizations and in those the bulla resonance is close to the fundamental frequency (Figs 5, 6). The courtship calls of other species are characterized by low fundamental frequency and rich content of spectral energy in higher harmonics. In those, the predicted resonance frequency of the bulla was close to one of the upper harmonics (Fig. 6, Supplemental Figs 1, 2). For the Common Goldeneye courtship call A we saw an intriguing overlap of each predicted bulla frequency with a separate peak in the power spectrum, while in courtship call B only the right bulla’s resonance frequency overlaps with a peak (Fig. 6H, I). In the Common Shelduck, we were only able to observe the left bulla overlapping with a courtship vocalization peak, while we were unable to match the predicted frequency of the right bulla with any vocalization peaks (Fig. 6R).

**Figure 5.**
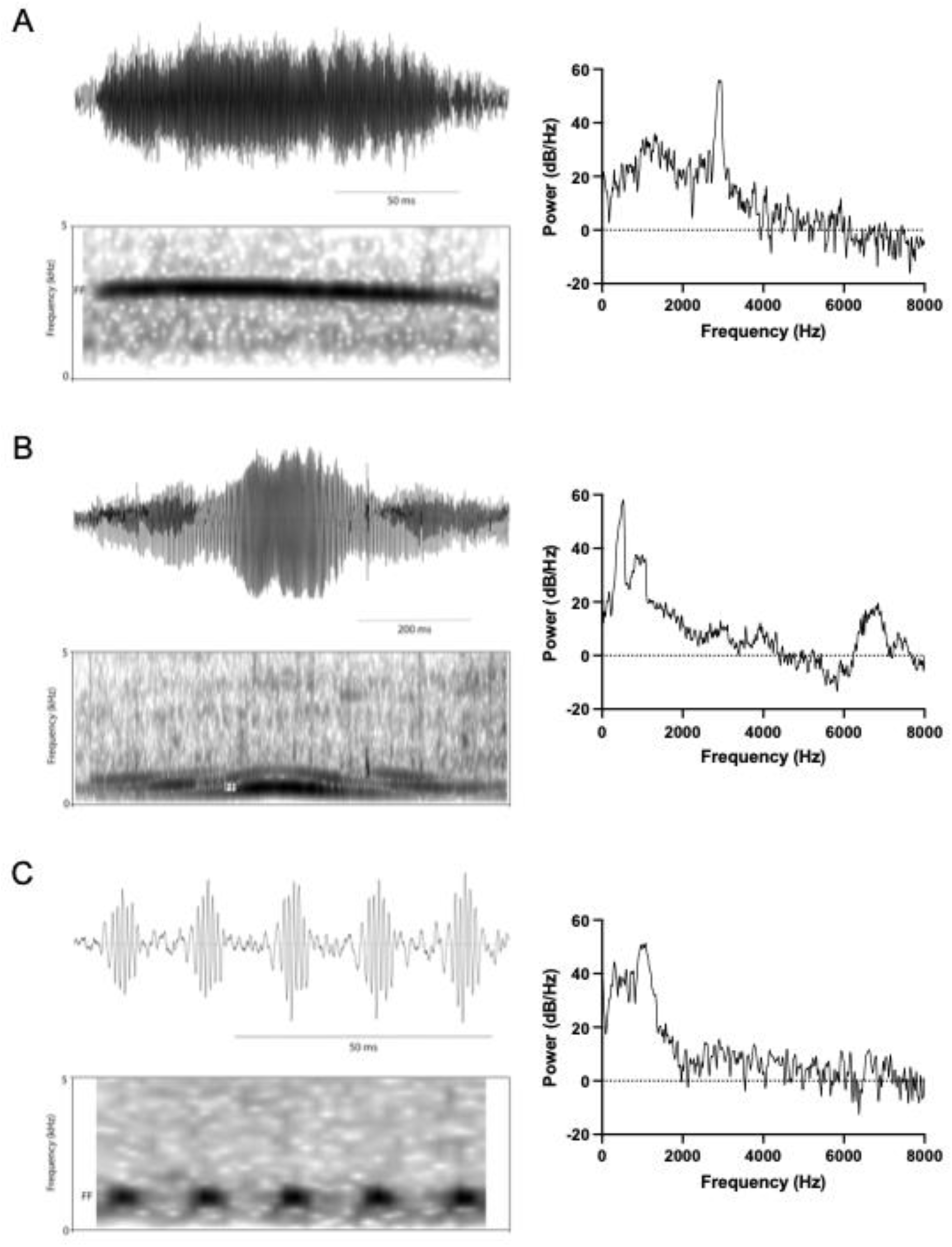
Courtship call examples. Left: above, oscillogram and below, spectrogram; right: power spectra of vocalizations to the left. (A) American Black Duck, which has a whistle-like vocalization (B) Common Eider, which has a low fundamental frequency and a complex call, (C) Hooded Merganser, which has a low fundamental frequency and a simple call.

**Figure 6.**
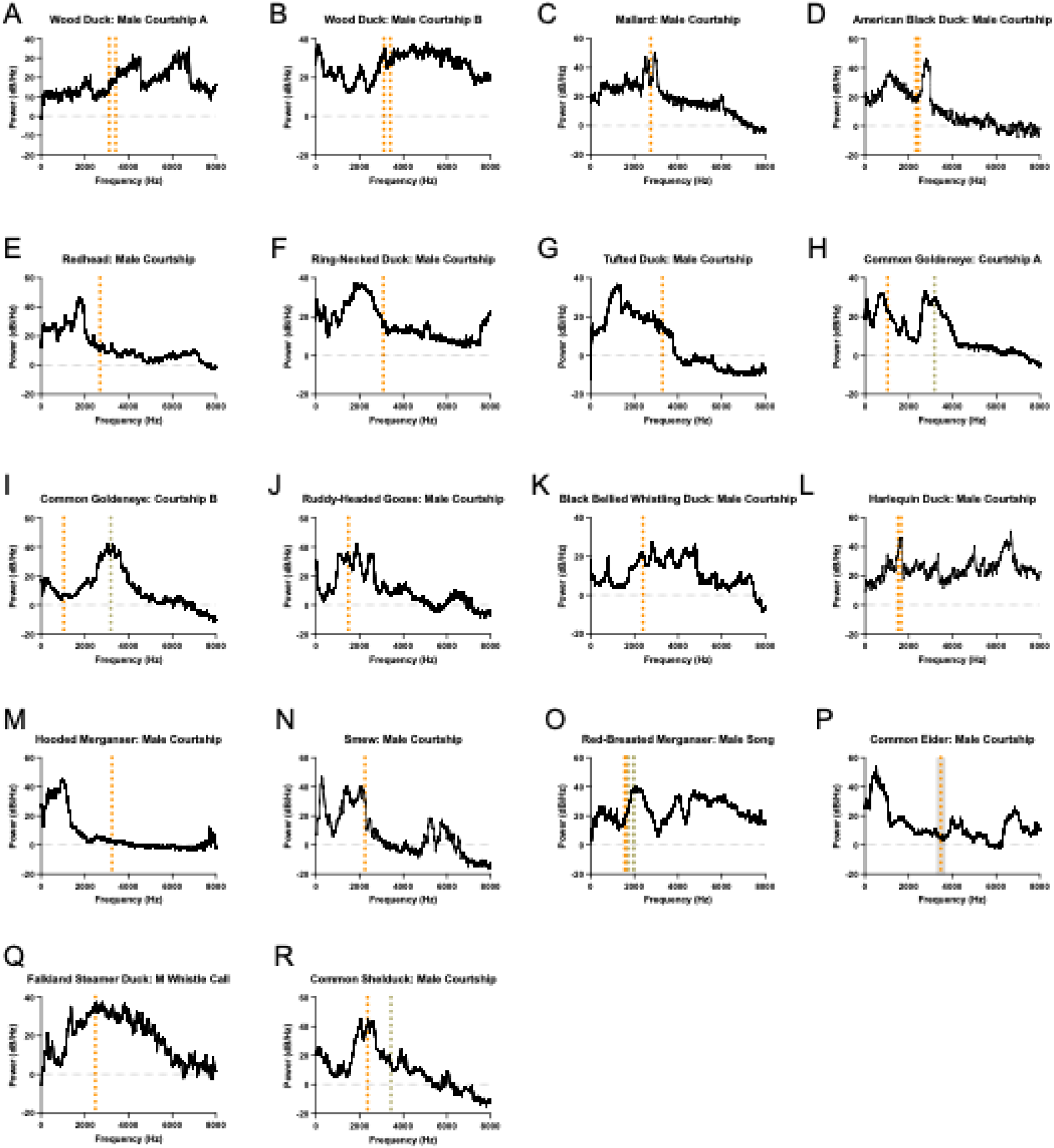
Courtship vocalization power spectra compared to predicted resonance frequencies of the bulla. Power spectra of one to five separate recordings were combined to produce power spectra for a given call type. Vertical orange dotted line represents the predicted resonance frequency of the left bulla. Vertical grey dotted line represents the predicted resonance frequency of the right bulla. Additional dotted lines indicate predictions from additional samples. Mallard is reproduced for comparison (Mishkind et al., 2024). Common Eider, the only sample to have five replicates, is shown with grey shading representing ± 1s.d.

Finally, for 3 species (Tufted Duck, Hooded Merganser, and Common Eider), we could not find a match between the spectral peaks of call types and the predicted resonance of the bulla. Tufted Duck has a complex courtship call that when taken as a whole shows an initial peak followed by an irregular region at around 20 dB below the peak (Fig. 6G). However, the fundamental frequency is modulated throughout the call, which leads to an unclear distribution of energy for the power spectra across entire calls (Fig. 7). A different story emerged when we looked at spectra at single points in the vocalization: there were some where the fundamental was emphasized, the second harmonic suppressed, and a more prominent third harmonic overlapped with our bulla prediction (Fig. 7).

**Figure 7.**
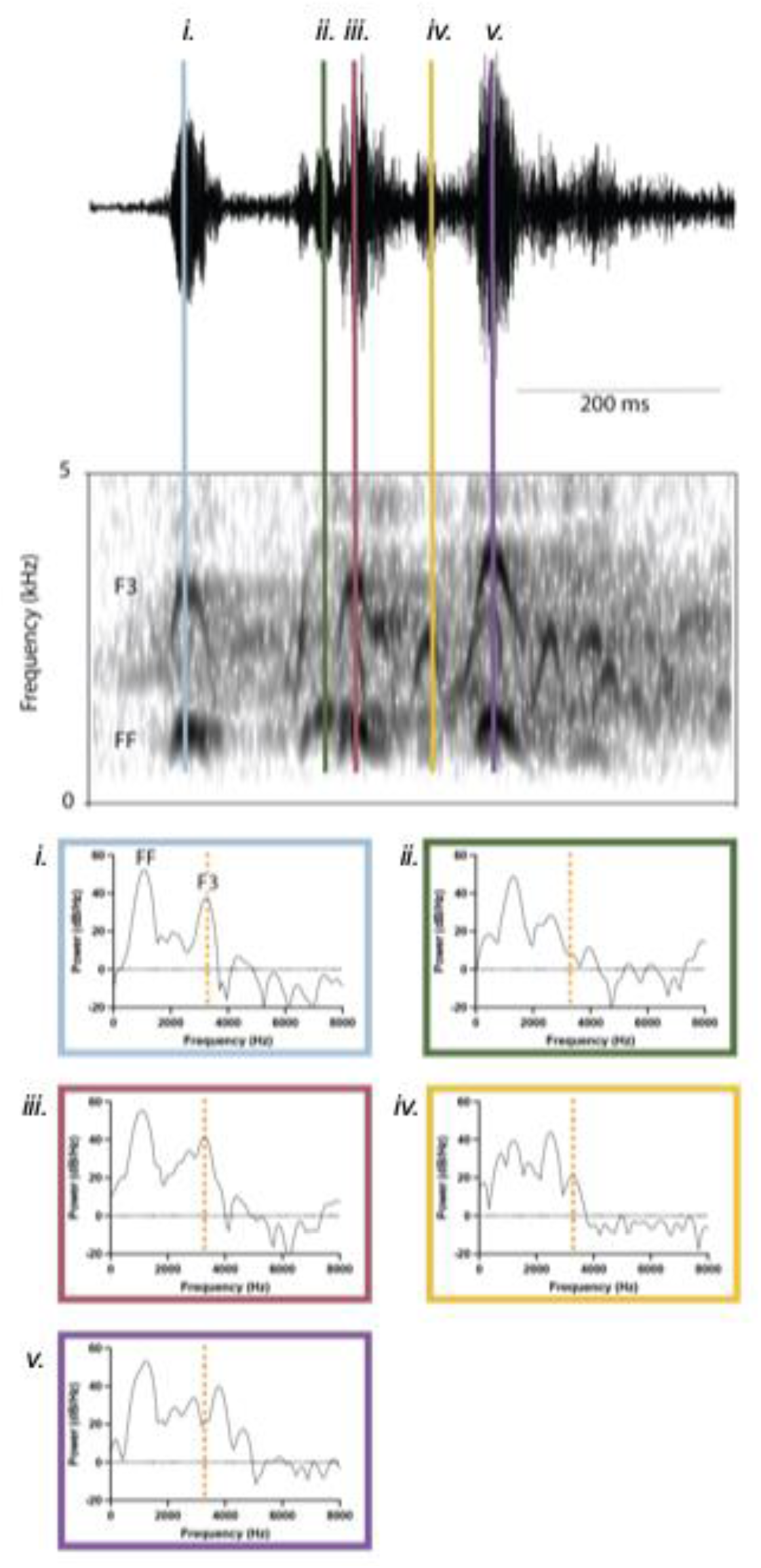
Tufted Duck courtship call is variable. Above, Tufted Duck courtship call shown as oscillogram and spectrogram. Below, power spectra of single points in the vocalization, locations spectra taken from indicated by colored lines corresponding to spectrum outline colors. Order left to right above corresponds to left to right and top to bottom in power spectra. FF, fundamental frequency; F3, third harmonic.

In the case of the Hooded Merganser, the low frequency courtship calls are likely generated in pulse tone mode with pulse repetition frequencies of approximately 55-65 Hz and each pulse consisting of a series of damped oscillations with a frequency of 1200-1600 Hz. Surprisingly for pulse tone vocalizations (Jensen et al., 2007; Sitt et al., 2008), the upper harmonic content in these calls is very low. It is unclear whether the low upper harmonic content is in part due to recording conditions (distance and noisy background) and/or reflects active suppression (Fig. 5C).

We then asked whether bulla volume and body mass showed a correlation. We hypothesized that outliers might represent species that had poor matches between our predicted bulla resonance frequencies and the recorded courtship calls. One striking outlier was the Common Eider, whose bulla is small relative to its body mass (Fig. 8). The Common Eider corresponds with our worst-matched estimate between courtship call and bulla resonance frequency (approximately 3 kHz difference; Fig. 6P). Overall, we observed a weak positive correlation between mean body mass and bulla size. When the Common Eider, Red-Breasted Merganser, Common Goldeneye, and Harlequin Duck were not included, we saw a regression line with a slope of 3.399 (p<0.0001) and an R^2^ of 0.9005 (Fig. 8). Two of the species that fell outside the general trend, the Red-Breasted Merganser and Common Goldeneye, both showed large bulla volumes for birds of their mean body mass. Interestingly, both species have left and right bullae. However, the third species we have studied that has both left and right bullae, the Common Shelduck, does fall within the general trend. The last species to fall somewhat outside the general trend, the Harlequin Duck, also has a moderately larger bulla volume than we see for others in relation to body mass.

**Figure 8.**
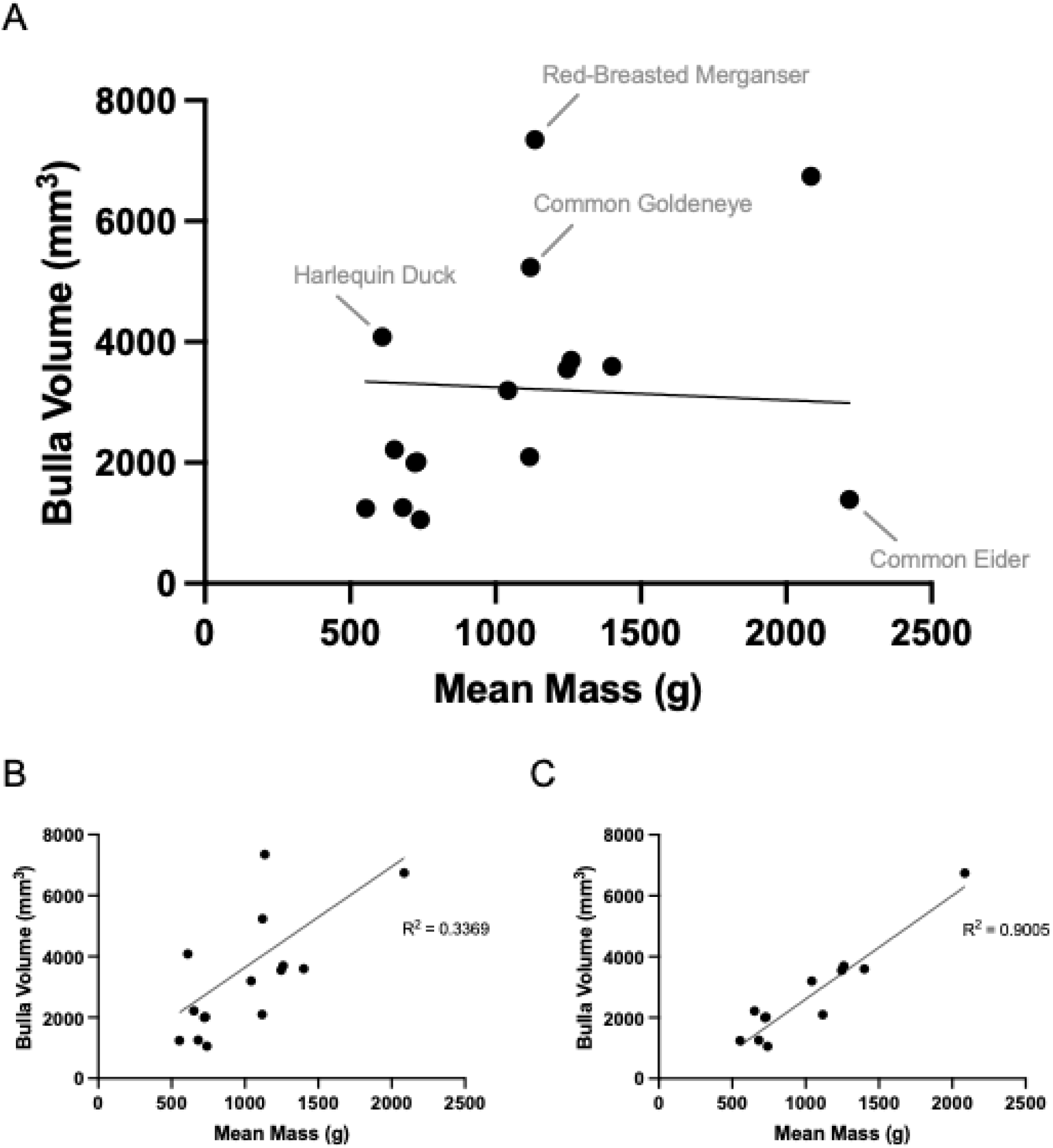
There is a weak correlation between bulla volume and mean species body mass. (A) Bulla volume vs. mean species body mass. Species outside the trend are labeled (R^2^ = 0.003, slope = −0.21, p value for slope = 0.78). (B) Plot with Common Eider removed (R^2^ = 0.34, slope =3.31, p value for slope = 0.0073). C. Plot with Common Eider, Harlequin Duck, Red-Breasted Merganser, and Common Goldeneye removed (R^2^ =0.90, slope = 3.4, p value for slope <0.0001).

Taken together, our results are consistent with the bulla functioning as a Helmholtz resonator, at least in most duck species. Moreover, our results support the idea that the bulla serves to influence resonance frequencies during male courtship vocalizations.

## Discussion

Despite much speculation about the possible roles of the duck bulla, its functional relevance has not previously been explored. Thus, we asked the specific question whether the bulla could function as a resonator to enhance courtship vocalizations. Predicted resonance frequencies for different species generally overlapped with peaks in spectral energy of courtship vocalizations. However, we also saw matching of predicted and observed resonance peaks in other male vocalizations and in female calls. While the male resonance peaks in non-courtship vocalizations can of course also be attributed to the bulla, in females, other resonances such as those of the trachea and OEC must account for the specific filtering. This latter conclusion therefore suggests that specific spectral properties of vocalizations arise from combined, and possibly overlapping, action of the various components of the upper vocal tract.

The trachea constitutes a tube whose resonance can be estimated. Assuming that ducks do not actively change tracheal length very much (Fitch, 1999; Daley and Goller, 2004; Ohms et al., 2010) we can use the range of measurements to calculate resonance frequencies. In the Mallard, tracheal length ranges between 14–18 cm (Klingler, 2016; Al-Ahmed and Sadoon, 2020), resulting in a resonance frequency of 476–613 Hz, if the trachea is seen as a tube with one open end. The fact that the fifth harmonic of the tracheal resonance frequency aligns with the predicted resonance frequency of the bulla may explain the overlap between our prediction and the peak frequency of the female Mallard flight call. This highlights one difficulty of evaluating the bulla predictions. After sound is produced by the syrinx, many components of the vocal tract beyond the bulla, including the trachea (Abs, 1969, 1970a), glottal opening, OEC and beak opening, influence which frequencies are enhanced (Fletcher et al., 2006; Riede et al., 2006; Kazemi et al., 2023).

The comparison between male and female Mallard ducklings provides a test to what degree the presence of a bulla in males alters the spectral distribution of energy in their vocalizations. We used contact (distress) calls, because these can be readily elicited by isolating the duckling and their context specificity allows direct comparison. Furthermore, they are uttered with high amplitude and are not contaminated by calls of other ducklings as would be the case in a social group. Surprisingly, there are no differences between calls from males and females with the exception of one-month-old birds. The 20 dB higher second harmonic of male calls could have arisen from a match with the resonance frequency of the bulla. Although we do not have measurements of the size of the bulla at this age, the 4–5 kHz resonance suggests a volume of approximately 540 mm^3^ (for 5 kHz) to 840 mm^3^ (for 4 kHz) assuming neck length and opening diameter are half the adult measurements. Conversely, we can ask why at all other times the bulla did not produce a match with harmonics in the calls. Given the bulla is still growing during this period, it may be a result of the bulla resonance frequency not matching the call frequencies until this time. Measurements of the growing bulla over time would be necessary to assess this.

Age was not included in sample annotations of our museum specimens, though research in male King Eiders and Common Eiders has shown that the size of the bulla did not vary between one-year olds and adults (Miller et al., 2007). Furthermore, in the species for which the most individuals could be sampled, the Common Eider, we only found small individual variation (coefficient of variation = 6.09%), which is consistent with published results in King and Common Eiders (Miller et al., 2007).

Our results constitute support for the hypothesis that the bulla of male ducks provides resonance properties that enhance aspects of their courtship vocalizations. We can only estimate the resonance frequencies because opening diameter and neck length of the opening cannot be unambiguously determined. In addition, the skeletal elements in these museum specimens do not give insight to what degree dynamic adjustments of the opening plays a role. For example, the close proximity of the labia and other soft tissue (Warner, 1971) at the opening could lead to a changing opening diameter and different driving pressures could even facilitate its dynamic modulation. In addition, precise shape (King, 1989; Frank et al., 2007), soft tissue partial partitions within the bulla (Warner, 1971; Lockner and Youngren, 1976; King, 1989), and fenestrations covered by elastic soft tissue, as are found in the bullae of *Aythya* species (Warner, 1971; King, 1989), will also affect the resonance properties.

The comparative approach includes species with very different spectral properties of courtship calls, whistle-like calls, broad-band, harmonically rich calls, and pulse tone calls. This broad range of spectral characteristics also leads to a potentially different role of bulla resonance across our duck species. While tuning to the whistle frequency will enhance the total amplitude of the courtship call, in harmonically rich courtship calls, bulla resonances can contribute to enhancing specific frequency bands in the harmonic content. Finally, bulla resonances could also act as filters of specific frequencies, as may be the case in the low-frequency pulse tone calls. The wealth of acoustic characteristics of courtship calls in the Anatidae points toward an interesting evolutionary relationship between the marked sexual dimorphism in syringeal structure and vocal behavior. Although we do not know which acoustic properties of courtship vocalizations are ancestral within Anatidae, it is possible that harmonically rich calls represent original courtship vocalizations. Whistled courtship calls are prominent in the genus *Anas* and may therefore be derived, which is consistent with their phylogenetic position within Anatidae (Sun et al., 2017). Resonance properties of the bulla can contribute to the broad range of vocal properties. The variation of bulla volume relative to body mass points toward interesting species in this context. However, it is also possible that the evolution of vocal features has outpaced changes to the morphological structure and therefore we are seeing evolution of syringeal morphology in progress.

Here we tested whether the bulla might act as a Helmholtz resonator, which could contribute specific resonances for enhancing particular frequencies of vocalizations arising from labial oscillations. Our comparative approach yields correlational evidence that largely supports this hypothesis. While such a function of the bulla is the most plausible, the present evidence cannot rule out some of the proposed alternatives. The main uncertainty stems from the fact that we have very little physiological data on the sound production mechanisms in ducks. For example, it is not clear which anatomical structures are the main sound generators. While some authors identify labia, others postulate various membranes as the primary sound sources (Rüppell, 1933; Warner, 1971; King, 1989). An additional unsubstantiated claim is that the bulla itself may generate sound as air flows past or through it. As a potential whistle mechanism Johnsgard proposed an aeolian whistle (Johnsgard, 1961, 1971). Aeolian whistles generate sound as vortices are shed when air flows around a cylinder (Chanaud, 1970), and it may therefore not be the most likely mechanism in light of the morphology of the male duck syrinx. Alternatively sound could be generated with whistle mechanism that resembles that of human whistling (Azola et al., 2018) or a flow-excited Helmholtz resonator, akin to an airstream flowing over the opening of a bottle. In this latter case, the resonance properties of the bulla, as calculated here, will predict the generated frequencies. While whistle mechanisms are clearly not the source of any vocalizations with complex harmonic spectra, one of them could give rise to the courtship whistles within the genus *Anas*. Future work will be needed to determine whether different production mechanisms give rise to different vocalizations within the call repertoires of ducks.

## Methods

### Duckling vocalizations

Mallard ducklings were ordered from a commercial breeder and kept in the laboratory in a pen equipped with an infrared heating lamp in one corner. They were fed *ad libitum* with commercial food for ducklings and were provided water at all times. Starting at the age of 4 days, individuals were isolated in a recording chamber whose walls were lined with acoustic foam, for audio recording of contact calls (sometimes also called isolation or distress calls) (Kear, 1968). Approximately 100-500 contact calls were recorded with an Audiotechnica AT3032 omnidirectional microphone at 44.1 kHz with Avisoft recorder. After the recording sessions, body mass was determined to the nearest 0.1 g. Recording sessions for each individual were repeated every week. We recorded calls from 4 males and 4 females for 7 weeks. After this period, the vocal repertoire developed into more diverse vocalizations, and birds no longer readily produced the contact calls when isolated. Sex was determined either post-mortem or when sex was revealed after molting into breeding plumage in the following spring.

From each recording we selected 15 loud calls that were free of background noise. We analyzed the sounds using Praat software (Boersma and Weenink, 2024). For each call we used the central segment to determine the mean fundamental frequency and the relative amplitudes of the fundamental (FF), the second (F2) and third (F3) harmonics from power spectra. To do this, we generated power spectra and extracted frequency and amplitude values.

### Comparative analysis

Recordings used for analysis were primarily collected from Cornell Lab of Ornithology’s Macaulay Library. Any additional recordings required were gathered from xeno-canto.org as noted in Table 2. One to five separate recordings of each call type were combined to generate power spectra in Praat Version 6.4.07 (March 17, 2024) (Boersma and Weenink, 2024).

**Table 2.**
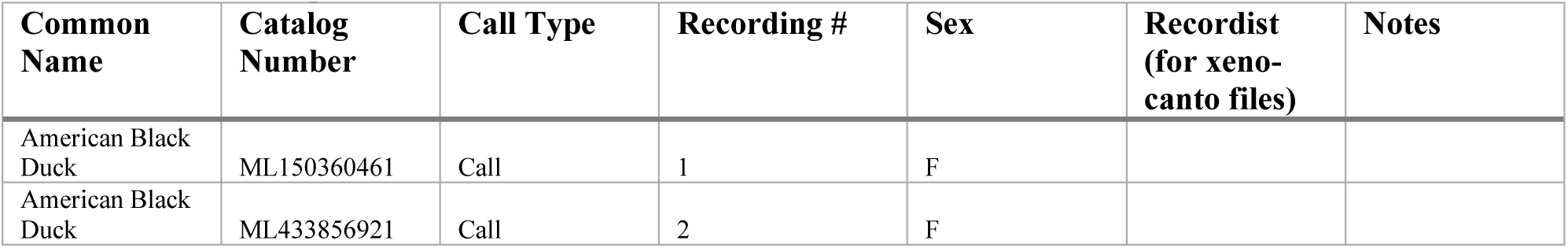
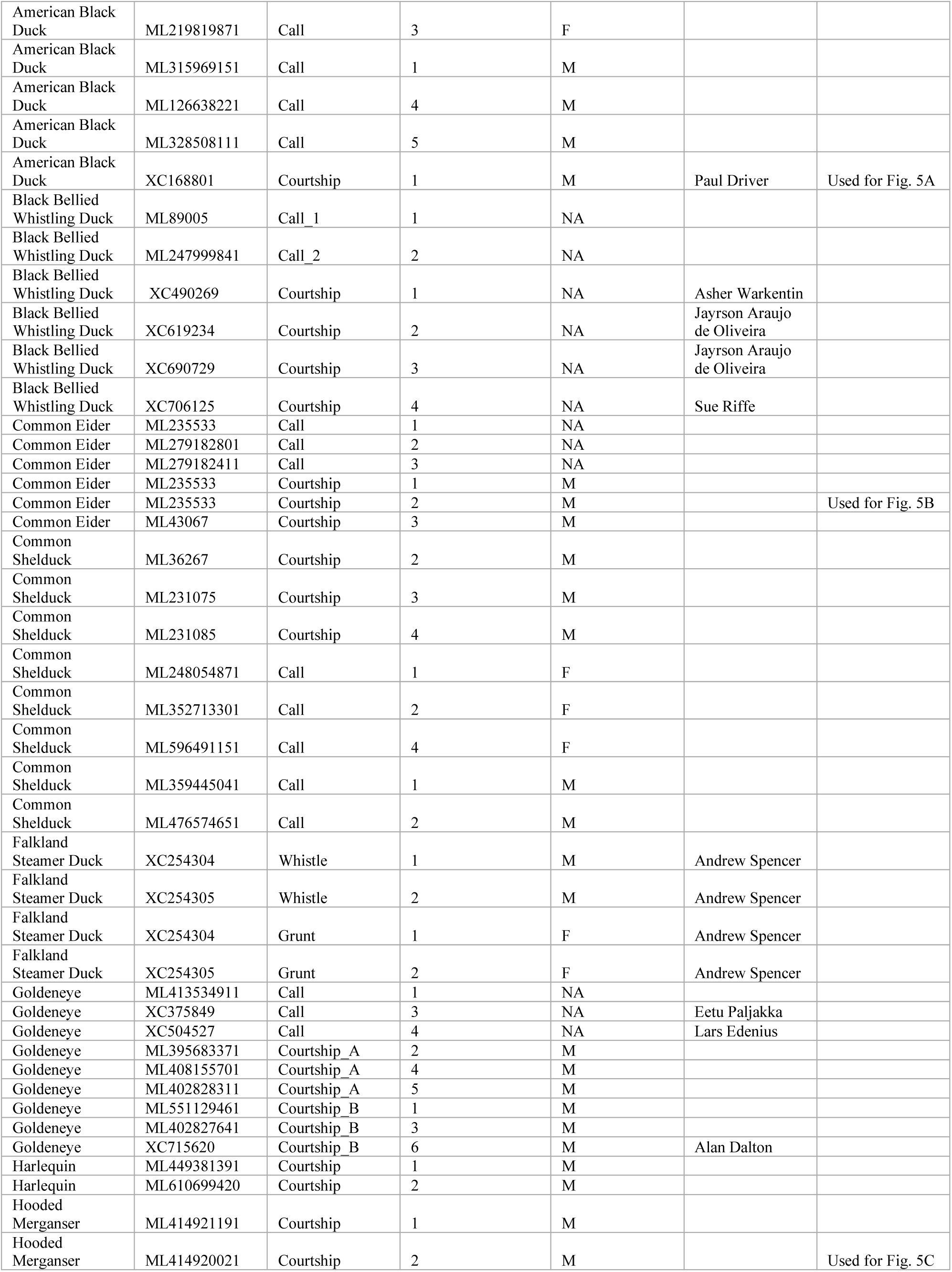
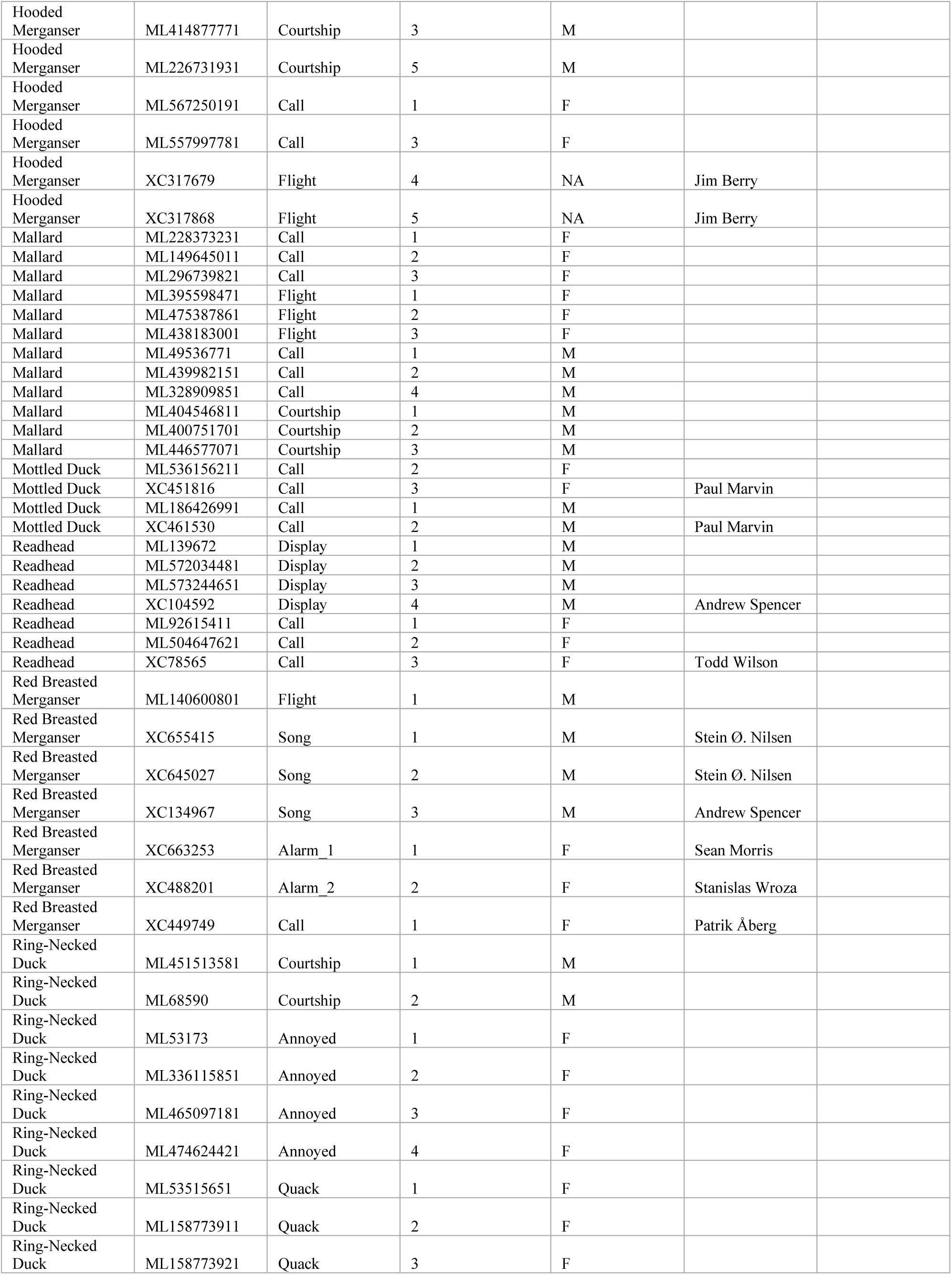
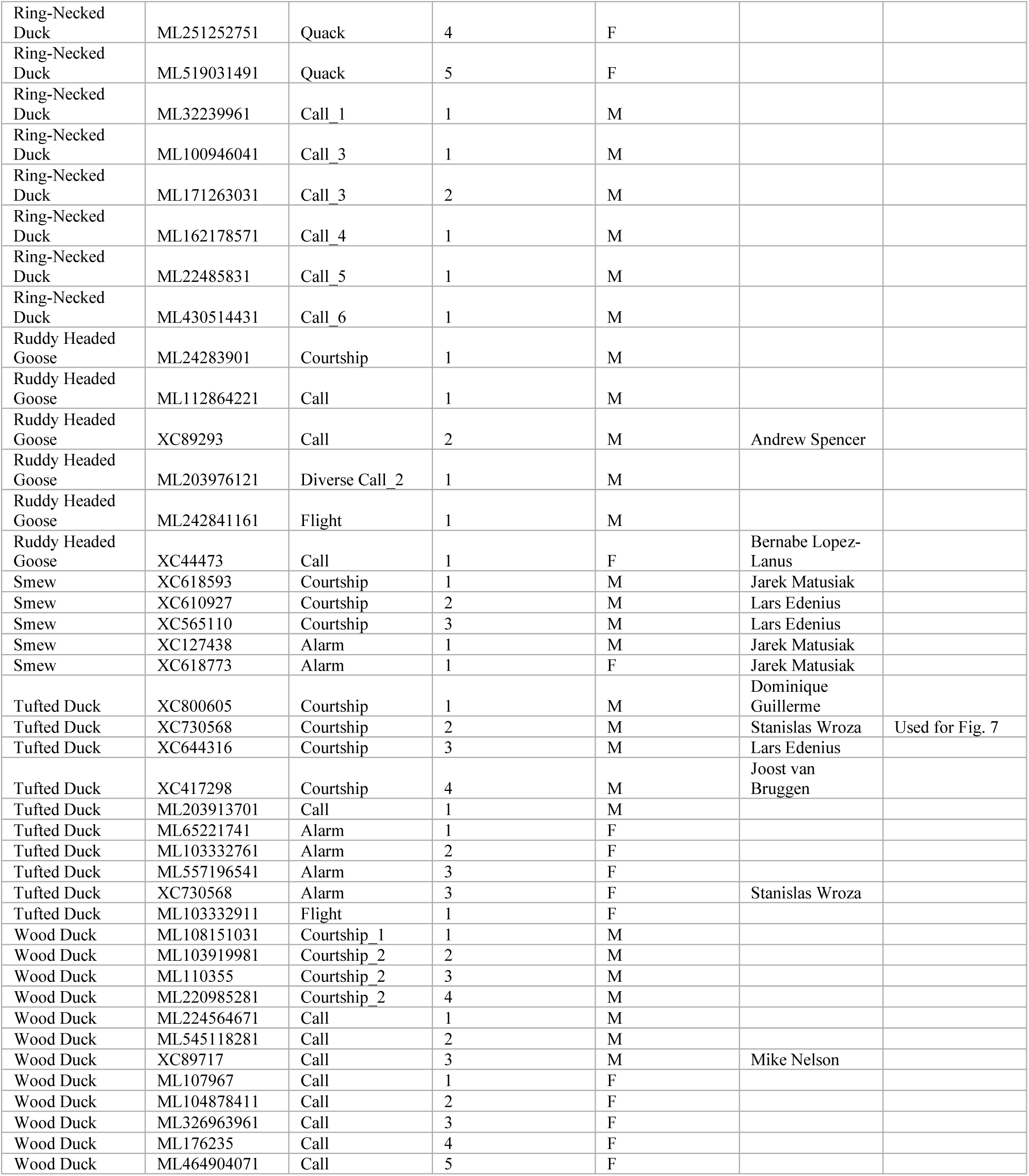
Sound files and their identifiers. Sound files used in figure 6 and supplemental figures 1 and 2, as well as in other figures as noted. For catalog numbers, the prefix ‘ML’ denotes that it came from Macaulay Library, ‘XC’ from xeno-canto.org.

Syrinx specimens were sourced from the Harvard Museum of Comparative Zoology ornithology department. Scanning with micro computed tomography (µCT) was conducted with X-Tek HMXST225 X-ray imaging system (Nikon Metrology, Inc., Brighton, MI, USA). Acceleration voltage was 60–75 kV and filament current 120–280 µA. Copper 0.1 filter was used for samples containing metal attachments: 342835 (Red-Breasted Merganser), 340383 (Smew), 340385 (Tufted Duck). 3142 projection images captured at 2000 pixels by 2000 pixels. Two frames were averaged with 1 second exposure time per frame. Pixel size = 0.2 mm.

Data reconstructed with Nikon CT Pro 3D software. Bulla volume, neck length, and opening diameter measured using Dragonfly Pro software, Version 2022.2 (Object Research Systems, Inc., Montréal, Canada). Measurements used to model bulla as a Helmholtz resonator:

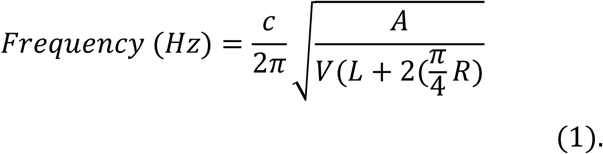

where c = speed of sound (343 m s^-1^), A = neck opening area, V = bulla volume, L = neck length, R = radius of neck opening area.

Literature indicated that several samples marked in the museum collections as female were, in fact, male (Bergmann and Düttmann 2001; British Ornithologists’ Union 1859; Miller et al. 2007; Wilson, Sonsthagen, and Franson 2013). Thus, we used a combination of museum annotations and literature to determine sample sex.

Graphing done with GraphPad Prism 10 for macOS Version 10.2.3 (347), April 21, 2024.

This work was performed in part at the Harvard University Center for Nanoscale Systems (CNS); a member of the National Nanotechnology Coordinated Infrastructure Network (NNCI), which is supported by the National Science Foundation under NSF award no. ECCS-2025158.

## Acknowledgements

We thank Jeremiah Trimble and Kate Eldridge of the Harvard Museum of Comparative Zoology Ornithology Collections. Hao-Yu Greg Lin at the Harvard Center for Nanoscale Systems for help with µCT and Natalia Verzhbitskiy for assistance with raising the ducklings.

