## Supplemental Figures for "Courtship vocalizations in male ducks: spectral composition and resonance of the syringeal bulla"

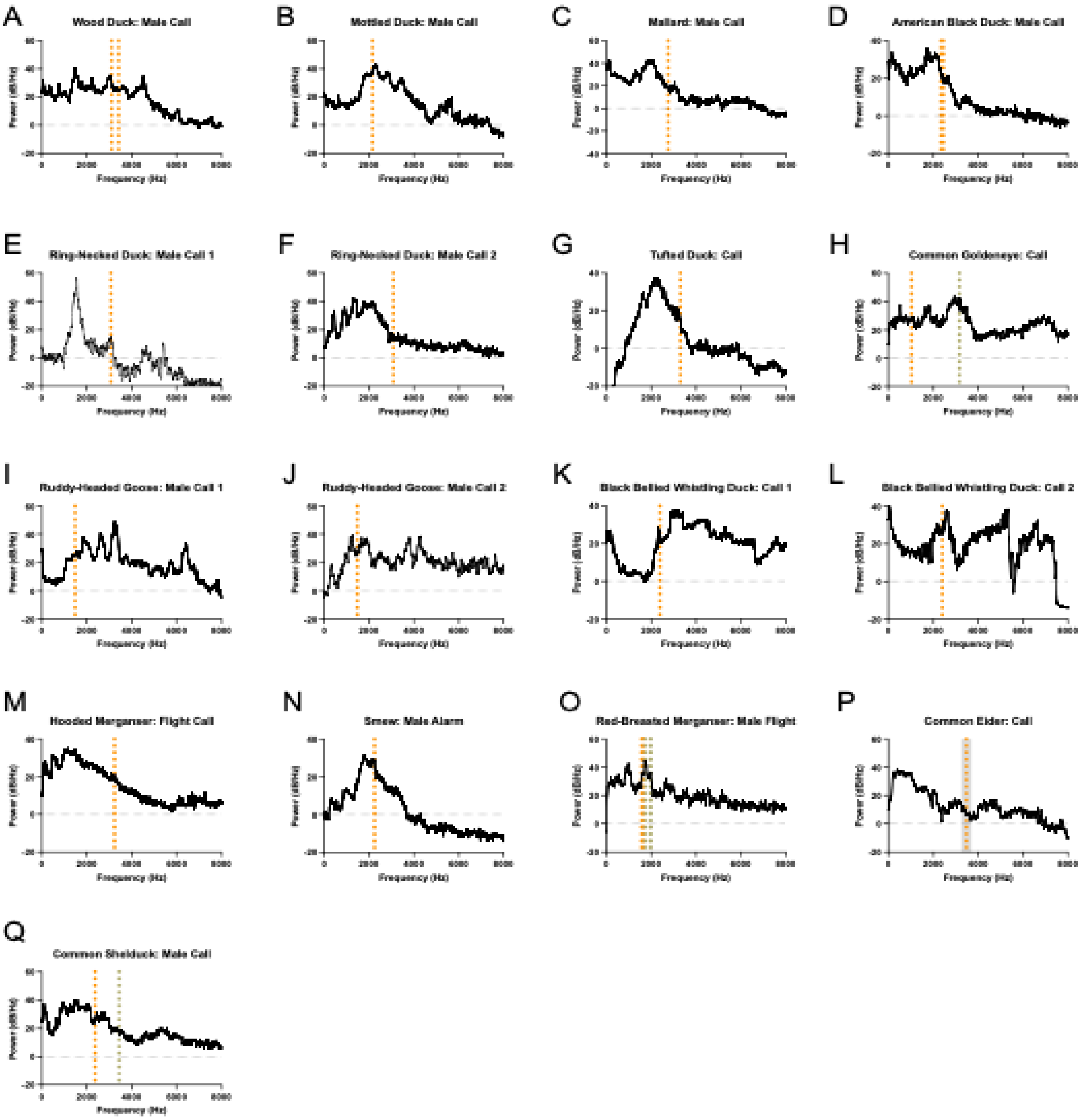


**Figure S1. Male non-courtship vocalization power spectra compared to predicted resonance frequencies of the bulla.** Power spectra of one to five separate recordings were combined to produce power spectra for a given call type. Vertical orange dotted line represents the predicted resonance frequency of the left bulla. Vertical grey dotted line represents the predicted resonance frequency of the right bulla. Additional dotted lines indicate predictions from additional samples. Mallard is reproduced for comparison (Mishkind et al., 2024). Common eider, the only sample to have five replicates, is shown with grey shading representing ± 1s.d..

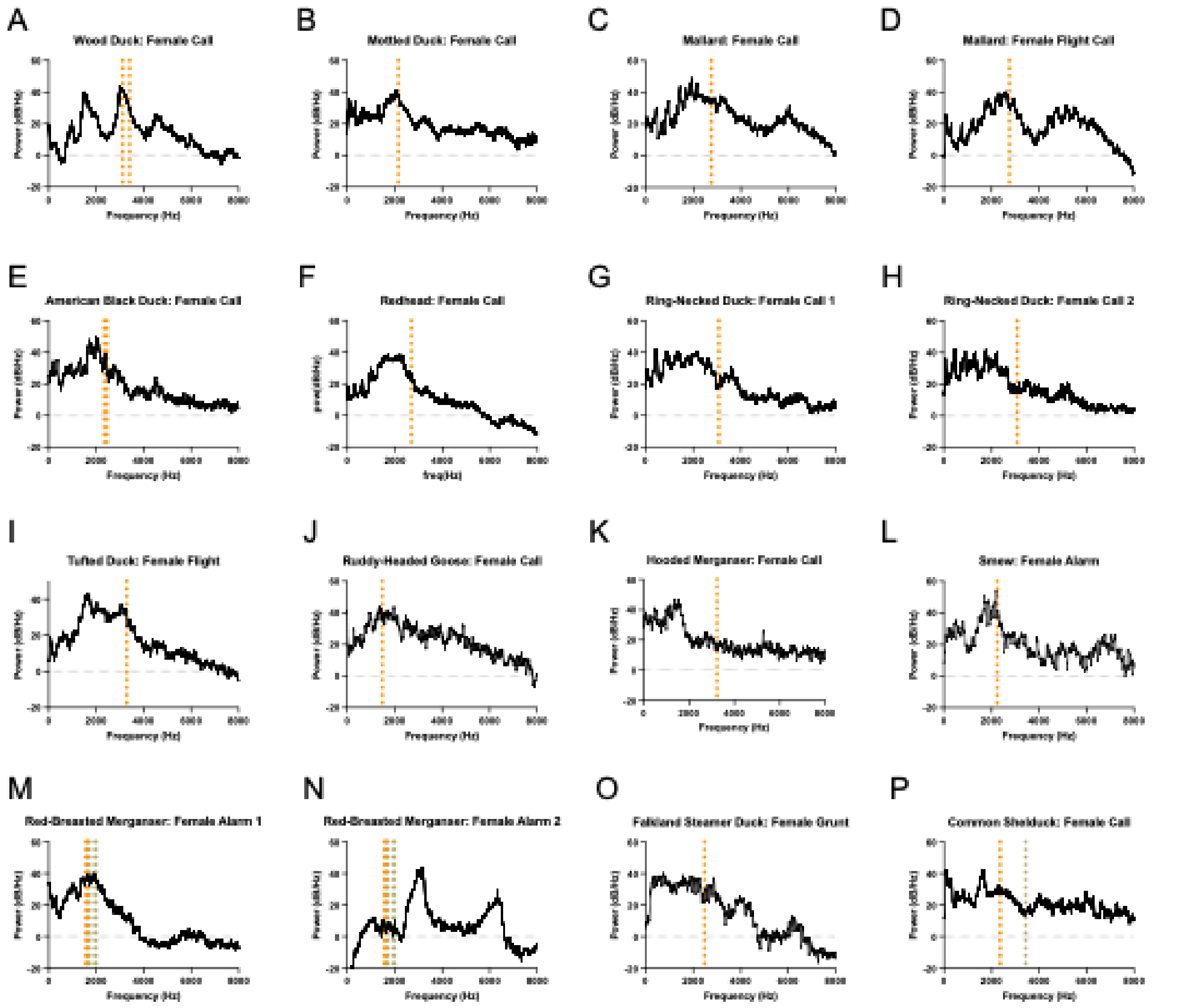


**Figure S2. Power spectra of female vocalizations compared to predicted resonance frequencies of the bulla.** Power spectra of one to five separate recordings were combined to produce power spectra for a given call type. Vertical orange dotted line represents the predicted resonance frequency of the left bulla of conspecific males. Vertical grey dotted line represents the predicted resonance frequency of the right bulla. Additional dotted lines indicate predictions from additional samples. Bulla resonance in the Common Eider, for which five male bullae were sampled, is shown with grey shading representing ± 1s.d..
